# Explainable AI shows climate impacts on wheat yields: insights from 30 years of field data

**DOI:** 10.1101/2025.09.19.677331

**Authors:** Margot Visse-Mansiaux, Masahiro Ryo, Amanda Burton, Tasin Siraj, Josepha Schiller, Simon Treier, Didier Pellet, Laura Stefan, Lilia Levy Häner, Juan M. Herrera, Thomas Cherico Wanger

## Abstract

Wheat (*Triticum aestivum L.*) is amongst the world’s most important staple crops and primary food source for an estimated 35% of the global population. Climate impacts have caused global yield stagnation and quantifying the climatic variables influencing wheat yield is critical to anticipate yield losses and design climate-resilient agricultural strategies. Here, we use a unique 30-year dataset on winter wheat variety trials in six sites across Switzerland, explainable artificial intelligence (XAI) and interpretable machine learning (IML) methods (*i.e.*, decision trees and gradient boosting models combined with post hoc tests) to elucidate climate drivers on wheat yields. We showed based on 405 varieties and over 10,000 observations, that climatic variables such as cumulative solar radiation, precipitation from sowing to harvest and genotype makeup are significant yield drivers. Partial dependence plots and variable interaction analyses revealed, for example, a yield plateau above cumulative solar radiation levels of ∼3000 MJ m⁻², suggesting complex genotype-by-environment interactions. These findings suggest that XAI adds important biological interpretability to predictive performance, and reveals the mechanisms how climate affects wheat yields. Our methodological framework and results can inform breeding activities, agronomic management, and adaptation strategies under climate change across environmental conditions in Switzerland and with global ramifications.

## Introduction

Wheat (*Triticum aestivum L.*) is one of the world’s most important staple crops and the primary food source for an estimated 35% of the global population^1^. In 2022, global wheat production reached 808 million tonnes second only to maize ^2^. According to the United Nations, wheat plays a central role in global diets: nearly 700 million tonnes are consumed annually worldwide. Wheat alone provides over 20% of the world’s calories and protein^3^ and contributes significantly to the nutrition of billions of people globally. In Europe, wheat yields have experienced periods of stagnation over recent decades. Previous studies have shown that long-term temperature and precipitation trends since 1989 have reduced wheat yields in Europe by 2.5% and contributed to yield stagnation, with climate trends accounting for about 10% of the observed yield stagnation^1^. In Switzerland, wheat production reached 487,000 tonnes in 2022, with an average grain yield of 5.5 t ha^−1^, which is considerably higher than the global average of 3.7 t ha^−1^ (Supplementary Figure 1) ^2^. Wheat yields in Switzerland increased steadily until 1990 and have remained stable since then (Supplementary Figure 1). However, yield stability varies amongst varietal genotypes, is constrained by environmental conditions^4^ and experiences considerable interannual variability by local weather conditions^2^ (Supplementary Figure 1). In the general context of climate change impacts on wheat production ^5,6^, Swiss bread wheat varieties exhibit significant differences in yield and quality stability depending on local soil and climatic conditions^7^. Moreover, Swiss winter wheat varieties differ in their exposure to heat stress under future climate scenarios, with earlier genotypes partially escaping heat events by reaching sensitive stages sooner in the season^8^. Yet, even early varieties may face increased heat stress in the coming decades, highlighting the need for further breeding strategies combining phenological adaptation and heat tolerance traits^8^. This need is reinforced as climate change is already affecting crop production in Europe, especially in Southern regions, where warming combined with drying trends has led to yield declines of 5% or more since 1989, as observed in Italy^9^. Experimental trials, showed that extreme weather events including heatwaves, droughts, excessive rainfall, and low solar radiation, led to significant yield penalties in most European wheat cultivars, although some varieties from specific regions exhibited better tolerance^10^. Most of these studies have used crop modelling and simulations^10–14^ but the use of new machine learning techniques that can be applied to scarce long term datasets remains limited.

Switzerland offers an ideal case study for understanding wheat responses to climate variability with new machine learning approaches due to its representative ecoenvironmental zones and availability of unique long-term wheat dataset. First, the country in the center of Europe spans a wide range of the continent’s climatic zones and elevations, making it a relevant test case for assessing climate impacts on wheat production. Second, compared to simulation experiments, historical records provide empirical evidence of how real-world factors, such as varieties, affect yields under actual climate conditions. While models can incorporate these factors, they often require long-term, crop-specific field data for calibration. Such long term datasets for wheat^4^, are valuable for statistical and machine learning (ML) approaches to disentangle climate effects on yield. ML techniques can outperform traditional models and provide a better predictability by handling nonlinearities and interactions^15,16^, but complex models like ensembles or deep learning are often viewed as black boxes. Explainable artificial intelligence (XAI) and interpretable machine learning (IML) aim to improve model transparency by revealing how predictions are made^17–19^. These tools can help to i) identify the relative importance of predictor variables on yields (permutation variable importance) ^20,21^, ii) explore predictor variable interactions^22^, and visualize variable association between predictor and yield (partial dependence plots)^23,24^. Applying IML to long-term wheat data in Switzerland will reveal complex genotype-by-climate interactions and offer further insights for climate-resilient wheat breeding in Switzerland but also as a model for temperate cropping systems worldwide.

Here, we use a unique 30-year dataset on winter wheat variety trials in six sites across Switzerland and explainable artificial intelligence (XAI) and interpretable machine learning (IML) methods (*i.e.*, decision trees and gradient boosting models combined with post hoc tests such as partial dependence plots and variable importance measures) to elucidate climate drivers on wheat yields. We show based on 405 varieties and over 10,000 observations, that climatic variables such as cumulative solar radiation, precipitation from sowing to harvest and genotype makeup are significant yield drivers.

## Results and discussion

### Using XAI methods to understand climate effects on wheat yield

We performed a descriptive analysis on the 30 years of Swiss national variety trial data, comprising over 10,000 wheat yield observations across 405 varieties and six sites. Grain yield showed no temporal trend, ensuring data stationarity for ML analysis. ANOVA revealed that “variety” consistently explained the largest share of yield variability, while the influence of “year” and “year x site” interactions increased over time. The findings indicate a growing impact of climate variability on yield, supporting the inclusion of climatic predictors in ML models. For details see Supplementary text S1.

To disentangle the effects of genotype and climate on yield, we applied two IML models, *i.e.*, DT and GB. Then we used different XAI methods to assess the contribution of predictors. The DT and GB models were performed on the train dataset [N = 8756] using 5-fold cross resampling with different tested sample sizes (*i.e.,* 7005, 7005, 7005, 7005 and 7004). For each dataset (*i.e.,* 1, 33, or 34 predictors), the best models were deemed those with the smallest RMSE. Relevant model values are given in Supplementary Table 6.

The test dataset [2162] was used to assess model performance and make predictions across three datasets. Figure 1 compares R-squared values from correlations (Pearson) between observed and predicted values for DT and GB models using only variety (Figure 1A), variety with climatic variables and the duration of the three periods (Figure 1B), and only climatic variables and the duration of the three periods (Figure 1C) as predictors. Results show that the GB model captures variety-induced variability better, with increased prediction accuracy (R² = 0.74) when including variety compared to using only climatic variables (R² = 0.59). In contrast, the DT model’s prediction accuracy remains similar with or without variety inclusion (R² = 0.51 and 0.50, respectively). This highlights the GB model’s ability to leverage variety effects, aligning with previous study^23^, who notes DT’s limitations and GB’s robustness through ensemble learning.

**Figure 1.**
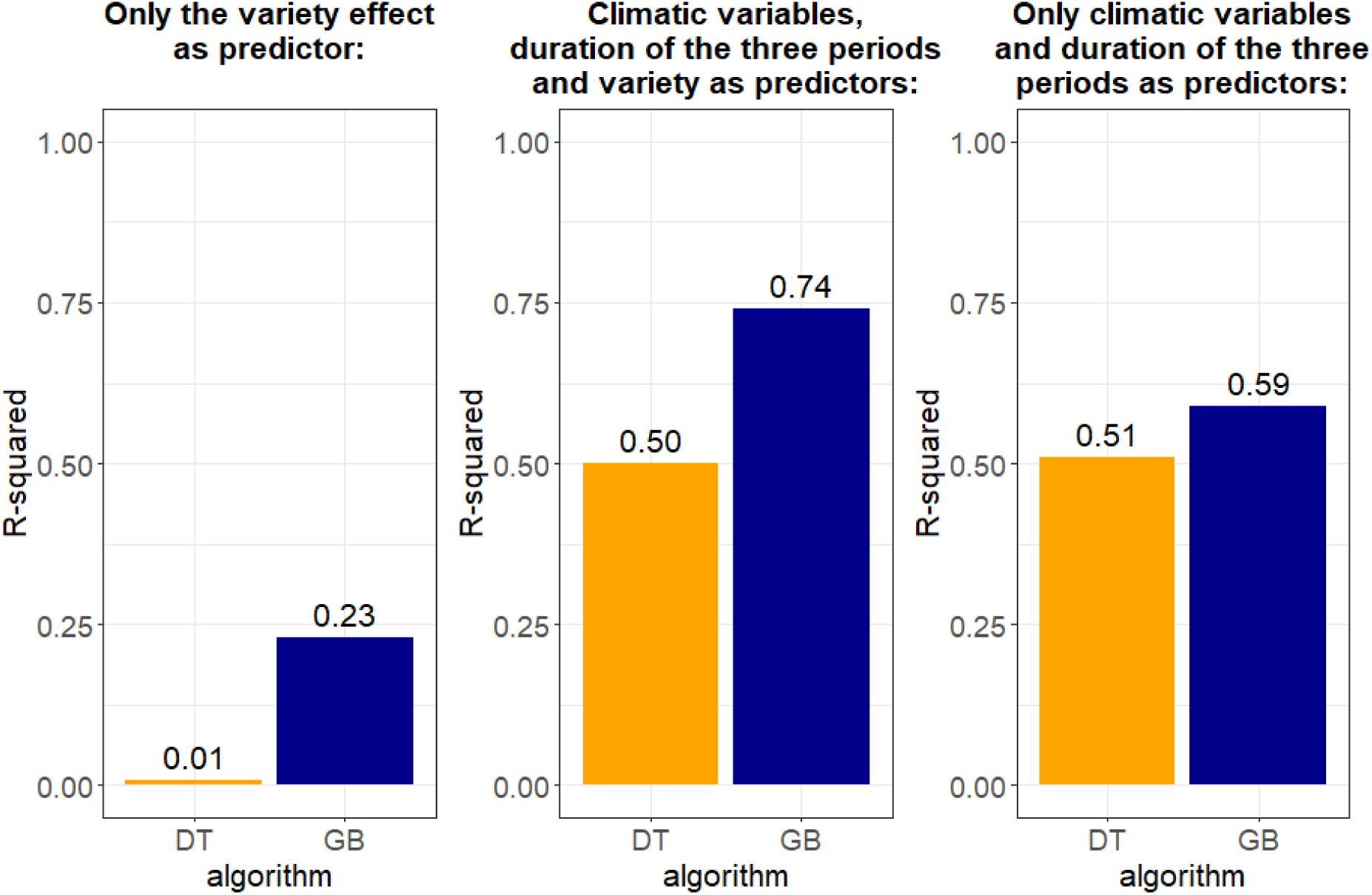
Comparison of R-squared values from the correlations between observed values and predicted values to evaluate the model performance of the two tested models: Decision Three (DT) and Gradient Boosting (GB) models performed with different predictors indicated on the graphics (*i.e.*, A - Only the variety effect as predictor of grain yield, B - With climatic variables, duration of the three periods and variety as predictors and C - Only climatic variables and duration of the three periods as predictors).

GB model consistently outperformed DT in predicting grain yield, even without variety effects (Figure 1B and C). Given the genotype’s consistency over decades (Supplementary Figure 5), subsequent analyses used only the dataset with 33 climatic predictors to focus on climatic impacts on grain yield. Variable importance for DT and GB models (Figure 2) identified key predictors: daily average precipitation (prec_p1), daily solar irradiance (ray_p1), minimum temperature during heading to harvest (tmin_p3), and cumulative solar radiation (somray_p1). Pairwise interactions among these variables were assessed using the feature importance ranking measure (FIRM), revealing strong interaction effects for both models. For GB, the strongest interactions occurred between prec_p1 × somray_p1, followed by ray_p1 × somray_p1 and prec_p1 × ray_p1 (Figure 3B). For DT, similar patterns emerged, with additional interactions involving tmin_p3 (Figure 3A). Both figures underscore the critical role of solar irradiance, daily radiation, temperature, and precipitation in grain yield predictions.

**Figure 2:**
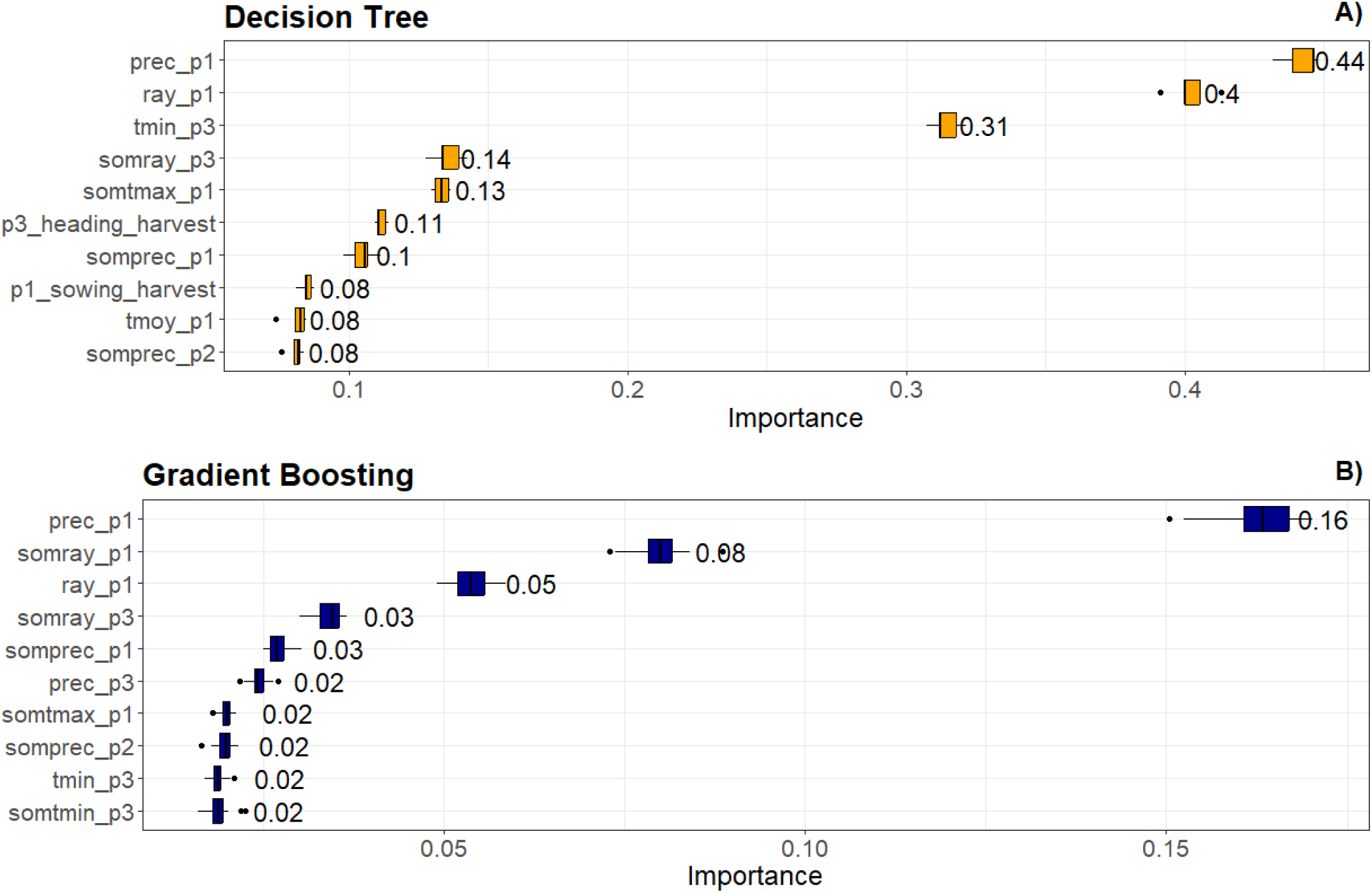
permutation-based variable importance (contribution to R-squared, ranging from 0 to 1) test showing the variable importance for the tested models, DT model (A) and GB (B). Legend: p1 = sowing to harvest period; p2 = sowing to heading; p3 = heading to harvest period; for each period: prec = average daily precipitation; ray = average daily irradiance; tmin = average daily minimum temperature; somray = cumulative solar radiation (sum of daily values x 24 x 0.0036 as there are 24 hours in a day and that 1Wh = 0.0036 MJ); tmax = average daily maximum temperature; tmoy = average daily mean temperature; somprec = cumulative precipitation (sum of daily precipitation); somtmin = cumulative minimum temperature (sum of daily minima); somtmax = cumulative maximum temperature (sum of daily maxima).

**Figure 3:**
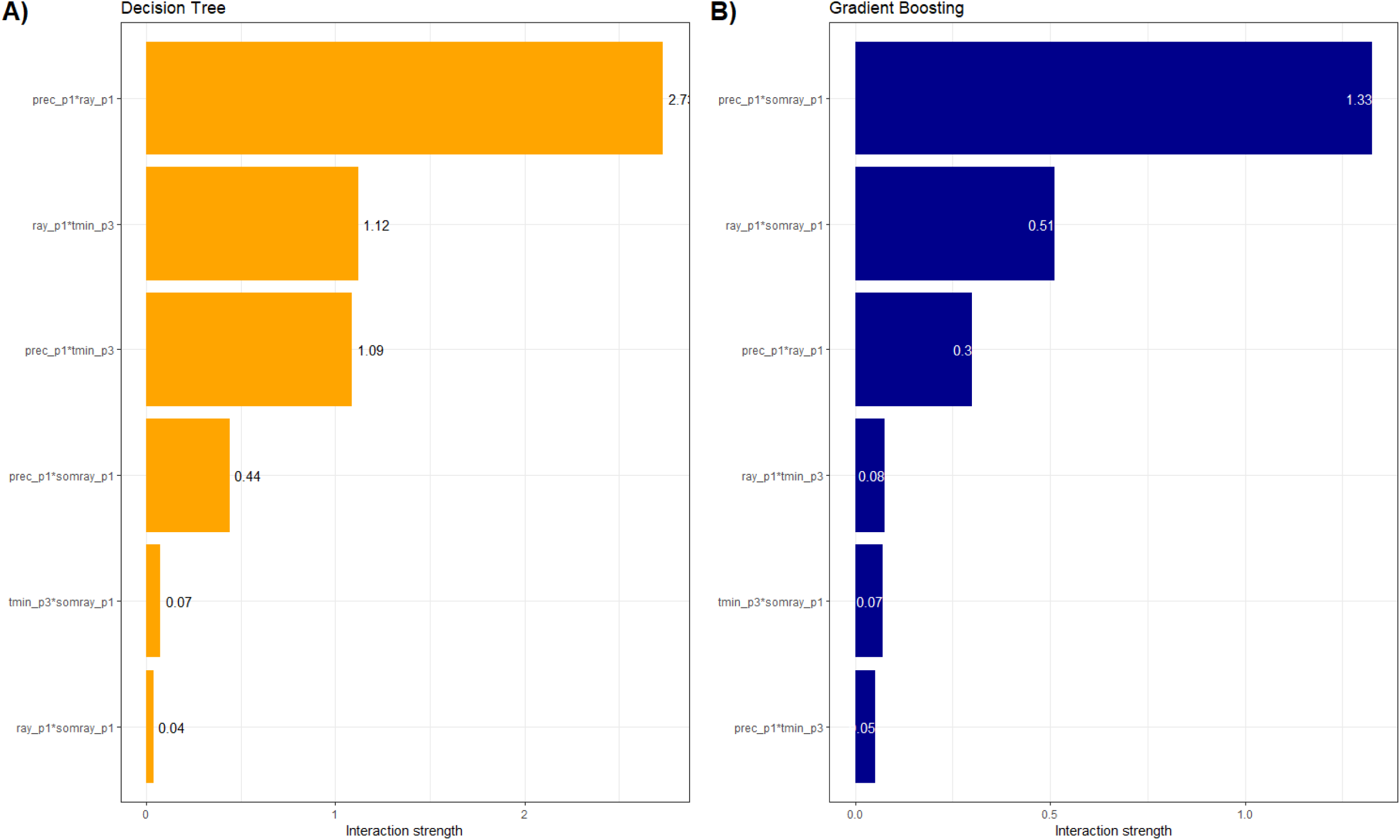
pairwise interaction importance (global model-agnostic method) based on feature importance ranking measure (FIRM) approach – Variable importance-based interaction statistic. This test was performed in the top three in variable importance tests by at least one algorithm for both models DT and GB. Legend: Legend: p1 = sowing to harvest period; p3 = heading to harvest period; for each period: prec = average daily precipitation; ray = average daily irradiance; tmin = average daily minimum temperature; somray = cumulative solar radiation (sum of daily values x 24 x 0.0036 as there are 24 hours in a day and that 1Wh = 0.0036 MJ).

Partial dependence tests (Figure 4) further explored the four key variables identified by both models. Both GB and DT models suggest a nonlinear negative relationship between average daily precipitation (prec_p1) and yield, with substantial decreases observed up to 3 mm/day before stabilizing at higher values (Figure 4-A). The GB model reveals threshold behavior for both average daily irradiance and cumulative solar radiation (Figure 4-B-C), with yield increasing sharply until cumulative solar irradiance reaches approximately 2,800 MJ m^−2^, followed by a plateau at approximately 3,000 MJ m^−2^ due to potential limiting factors like nutrients or photosynthetic saturation (Figure 4-B).

**Figure 4:**
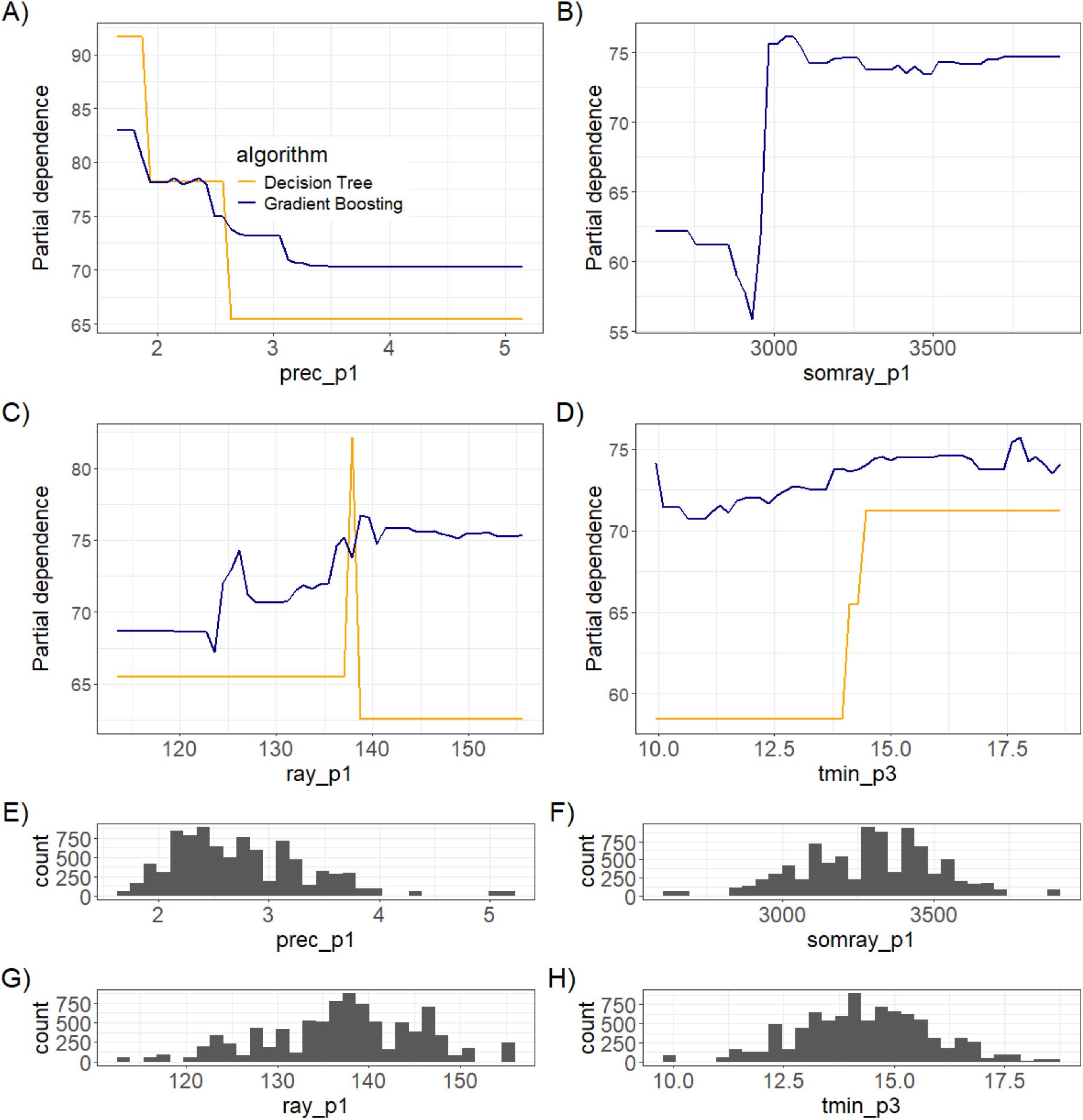
Partial dependence plots of the top three predictors identified as important in the variable importance tests by at least one algorithm for both models, DT and GB. Graphics A, B, C and D display the relationships between the predictors (*i.e.,* prec_p1, somray_p1, ray_p1, and tmin_p3) and grain yield, where the X-axis shows the values of each predictor and the Y-axis shows the partial dependence, corresponding to the average effect of the X variable on grain yield for each model (GB or DT). A flat curve indicates that the variable does not affect the prediction, whereas curves with a positive or negative slope indicate a positive or negative relationship between the predictor and grain yield. Graphics E, F, G, and H display the data distributions for these same variables. Legend: p1 = sowing to harvest period; p3 = heading to harvest period; for each period: prec = average daily precipitation; ray = average daily irradiance; tmin = average daily minimum temperature; somray = cumulative solar radiation (sum of daily values x 24 x 0.0036 as there are 24 hours in a day and that 1Wh = 0.0036 MJ).

Minimum temperatures during heading to harvest (tmin_p3) show a positive nonlinear relationship with yield for both models, with predicted yield increasing before reaching a plateau (Figure 4-D). Extreme agro-climatic factors, such as high temperatures, drought, and excessive rainfall, are supported in the literature as influencing yield variability across European wheat varieties^10^. These findings underscore the use of machine learning models in capturing complex agro-climatic interactions.

Figure 5 illustrates how raw key climatic factors (*i.e.,* prec_p1, somray_p1, ray_p1, and tmin_p3) influence wheat grain yield across the entire dataset, enabling a comparison with the trends observed in the post-hoc analyses based on the model outputs. These results suggest that, during the period from sowing to harvest, a global increase in precipitation tends to decrease yield, while higher levels of mean daily solar irradiance and cumulative solar radiation are associated with increased yields. We can also observe a threshold just below 3,000 MJ m^−2^ of cumulative solar radiation, with an increase in yield above this threshold, which is in line with results of partial dependence plots (Figure 4-B). Similarly, the decreasing trend of yield with increasing precipitation visible in Figure 5A corresponds well to the negative slope shown in the partial dependence plots (Figure 4A). For average solar irradiance(ray_p1), Figure 5C shows moderate positive trends that partly reflect the patterns observed in the partial dependence plots (Figure 4C), although the raw data exhibit more variability.

**Figure 5:**
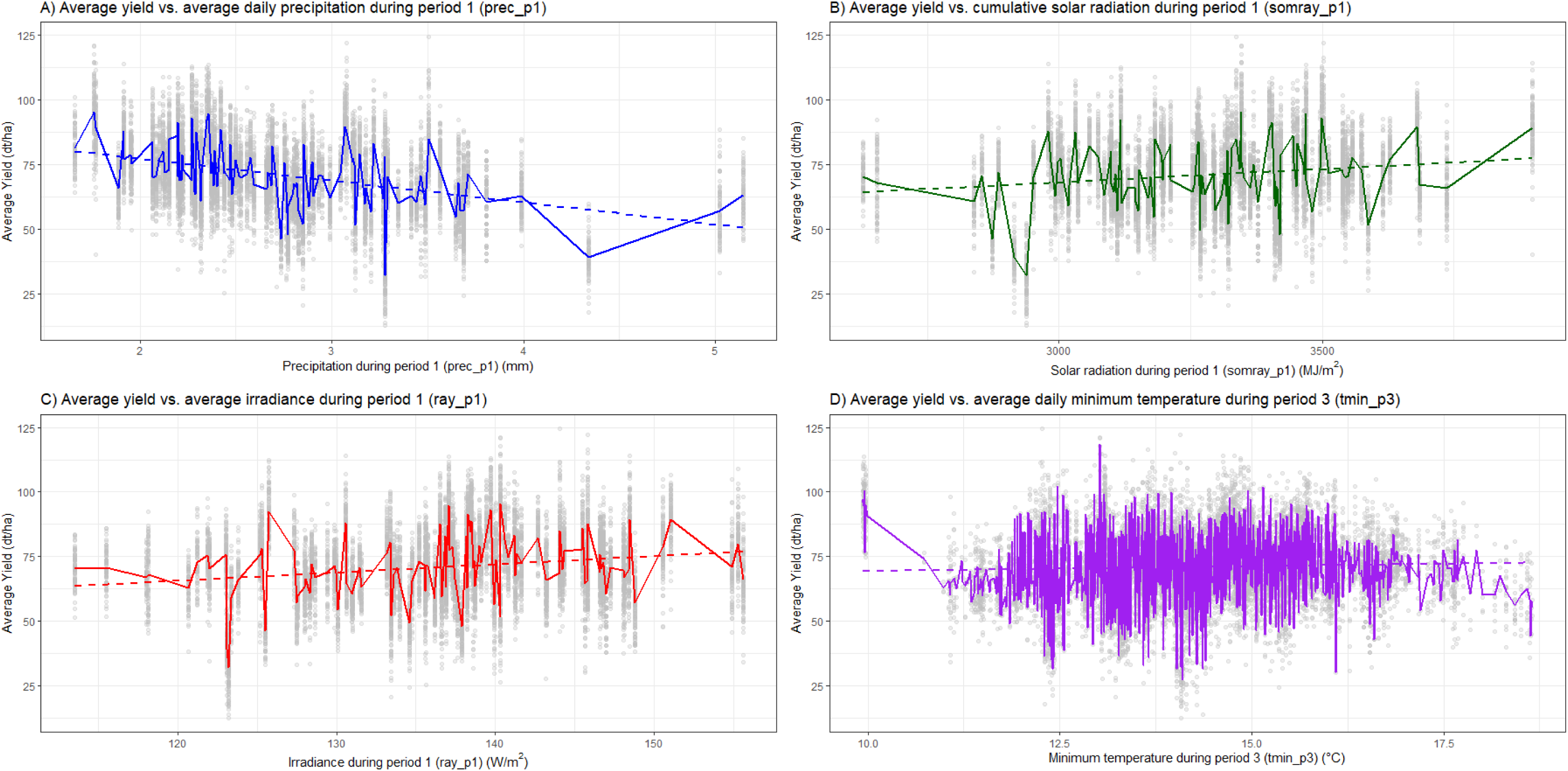
Impact of climatic factors on wheat grain yield for the entire dataset, showing the raw data points in grey, the smoothed curves (in colour) representing mean yield trends, and the linear trend lines (dashed lines) for A) average daily precipitation during the period from sowing to harvest (prec_p1), B) cumulative solar radiation during the period from sowing to harvest (somray_p1), C) average irradiance during the period from sowing to harvest (ray_p1), and D) average daily minimum temperature during the period from heading to harvest (tmin_p3).

In contrast, variations in minimum temperatures during the period from heading to harvest appear to have a milder effect on yield compared to precipitation, radiation and irradiance, although a slight positive trend is visible in both the raw data (Figure 5D) and the partial dependence plots (Figure 4D). It should be noted, however, that Figure 4 captures clearer thresholds and non-linear effects that may not be as easily visible in the raw data of Figure 5, highlighting the added value of model-based analyses.

Overall, this comparison confirms that the main relationships and thresholds identified by our machine learning models are also reflected in the observed data, supporting the reliability of the model outputs for understanding climate impacts on wheat yield.

Figure 6-A illustrates the variable importance of four predictors (prec_p1, somray_p1, ray_p1, and tmin_p3) from permutation-based feature importance tests using GB models on subsets for the 15 most tested wheat varieties. It compares the importance of these variables for individual varieties to their importance across the entire dataset. The results indicate that somray_p1, ray_p1, and prec_p1 are generally important for all varieties, but less so for popular varieties. Conversely, tmin_p3 appears to be more influential for popular varieties than for the overall dataset. This suggests that popular varieties may be less sensitive to climatic variability captured by somray_p1, ray_p1, and prec_p1 but are more responsive to tmin_p3. This interpretation aligns with Figure 4-D, showing a positive non-linear relationship between grain yield and tmin_p3, and effect amplified for popular varieties.

**Figure 6:**
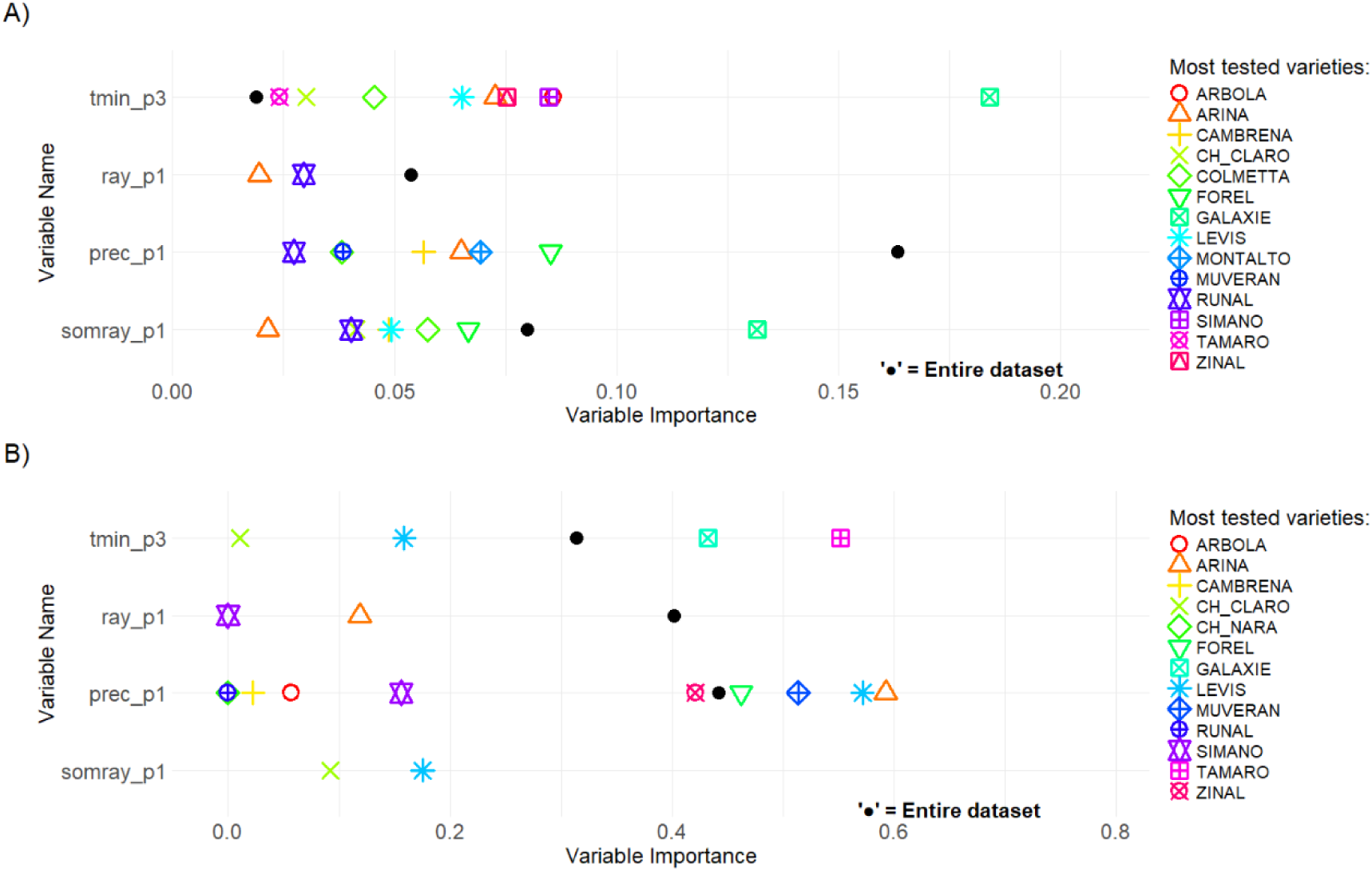
graph representing the variable importance of four variables (*i.e.,* top three variables in variable importance tests by at least one algorithm for both models DT and GB: prec_p1, somray_p1, ray_p1 and tmin_p3) from the permutation-based feature importance test on models performed for each variety among the 15 most tested varieties (shape and color for each variety in the legend) compared the variable importance from the permutation-based feature importance test performed on the model with the entire dataset representing the importance for all varieties (test on the entire dataset without the variety effect) (black dot), for GB model (A) and DT model (B). Among the 15 most tested varieties, if varieties are missing, it is because for the variety none of the four variables were present in the top ten most important variables in the variable importance test. Legend: p1 = sowing to harvest period; p3 = heading to harvest period; for each period: prec = average daily precipitation; ray = average daily irradiance; tmin = average daily minimum temperature; somray = cumulative solar radiation (sum of daily values x 24 x 0.0036 as there are 24 hours in a day and that 1Wh = 0.0036 MJ).

Figure 6-B presents the variable importance of the same predictors (except for somray_p1) using DT models on the same subset of varieties. The results for ray_p1 mirror those in Figure 6-A, confirming its significance across most varieties but not for the popular ones. For prec_p1, the DT model highlights its higher importance for specific varieties like Forel, Muveran, Levis, and Arina, compared to the entire dataset. This contrasts with the GB model where prec_p1 was consistently important across all varieties. The significance of tmin_p3 also varies by model. According to the GB model, tmin_p3 is more influential for popular varieties, while the DT model shows that only certain popular varieties (*e.g.,* Galaxie and Tamaro) assign greater importance to this variable compared to the overall dataset. These differences likely arise from model-specific methodologies and the relative weight given to each predictor. For example, prec_p1 is the most important variable in both models but shows a sharper distinction in importance in the GB model (Figure 2). In summary, while both models agree on certain predictors’ influence, differences exist in how correlated variables like prec_p1 and tmin_p3 are prioritized, reflecting the nuanced nature of these methods in capturing complex interactions.

### Climate impacts on Swiss wheat yields – Patterns and Benefits of XAI

In our study, yield predictions by models (Figure 4) and raw observed yields (Figure 5) decreased with higher daily average precipitation. Both daily radiation and cumulative solar irradiance from sowing to harvest significantly influenced yield predictions (Figures 2–4) and determined photosynthesis, impacting dry matter production and yield^25–28^. Observed yield patterns suggest initial limitations due to insufficient irradiance, sharp yield increases when cumulative irradiance reaches optimal levels at ∼ 3,000 MJ m⁻² because of photosynthetic saturation, and energy dissipation through non-photochemical quenching^29–32^. Our results are in line with other studies in Switzerland and Europe showing that wheat yields have been increasingly affected by adverse weather conditions such as excess rainfall, but also heat stress, and reduced solar radiation^4,10,33^, with significant genotype-by-environment interactions observed across different regions^7,8,10^. Wheat yield stagnation in Europe was attributed to climate trends^34^ while excess precipitation, for instance from China, led to yield losses and risks like disease and pre-harvest sprouting^35^. Our results demonstrate no temporal trend in mean grain yields in Switzerland since 1990 (Supplementary Figure 4-D); however, yields have shown considerable interannual variability driven by local weather conditions^2^ (Supplementary Figure 1). Minimum temperatures from heading to harvest period were associated with slight yield increases, consistent with findings that night warming enhances pre-anthesis growth and post-anthesis starch biosynthesis, boosting yield^36^. In contrast to most studies that assess climate variables independently to predict grain yield, our XAI framework enhances predictions by using ML combined with post hoc tests, including permutation-based interaction metrics. These tests revealed important two-way and synergistic effects, such as those between precipitation, solar radiation, irradiance, and temperature on grain yield (Figure 3). These findings highlight the added value of XAI tools, which allow not only accurate yield predictions but also interpretation of the underlying climate-yield relationships, revealing thresholds and nonlinear effects that would remain hidden in conventional statistical approaches.

#### Outlook

Realizing some of the limitations of this work are important future research avenues. First, the issue of handling highly correlated variables such as climate variables like somprec_p1 and prec_p1 is an often unresolved problem. While this correlation might suggest redundancy, these variables can represent different non-linear interactions and complementary information in crop responses (*e.g.,* daily precipitation vs. cumulative precipitation effects on yield), as seen in their differing variable importance rankings (Figure 2). Here we addressed this issue by retaining both variables in the model, based on their distinct agronomic interpretations, daily versus cumulative precipitation, and their differing importance rankings (Figure 2). While post hoc analyses focused on prec_p1 due to its higher predictive power, the inclusion of somprec_p1 ensured the model could capture potential cumulative effects of precipitation not reflected in daily averages. Correlated variables in ML therefore require further technical research attention. In addition, nonlinear ML models often outperform linear models when used with large datasets but they are less interpretable^37^. To address this, we used post hoc explainability analyses to quantify the impact of climatic variables on grain yield. This highlights the need for future research to improve explainability ML methods, especially for applications involving agronomic decision-making and breeding strategies.

Our analysis allowed us to identify key climatic constraints on winter wheat production in Switzerland, notably excess precipitation, while increases in irradiance and minimum temperatures had a positive effect on yield. On the one hand, these findings can guide future breeding activities as well as experimental studies, focusing on targeting identified climatic variables for different wheat varieties. This situation is particularly relevant for Switzerland, where frequent rainfall and limited solar radiation can pose major constraints on wheat yield stability^33,38^. In contrast, some other European and global wheat-producing regions face increasing risks from drought and heat^39^, making our findings complementary for guiding adaptation to region-specific climate conditions and breeding strategies. Overall, breeding efforts and new varieties should be integrated into diversified agricultural systems to advance a global food systems transformation^40–42^.

## Materials and methods

### 2.1) Data collection and preparation

The dataset was obtained from multi-environment winter wheat variety trials conducted at Agroscope, the Swiss agricultural research center in Switzerland for 29 years (from 1990 to 2020, without 1998 and 1999 due to missing data, see below). Trials were located in six different sites with elevations from 452 m [site “Changins” or “CH”] to 641 m [site “Grangeneuve” (GR)] (Supplementary Figure 2 and Table 1).

Sites were managed according to a Swiss regulation called “Principles of fertilisation of agricultural crops in Switzerland”^43^.

Grain yield data (*i.e.,* representing the dependent variable) was collected and recorded in the database, as well as the dates of the main crop stages (when available, *i.e.,* sowing date, harvest date and heading date). The variables: main stages, variety name, sites and year of testing were included in the database. Three periods (in days) based on these stages were calculated: 1) the period from sowing to harvest (p1); 2) the period from sowing to heading (p2); and 3) the period from heading to harvest (p3). Grain yield was measured on three replicates per variety, site and year.

Daily weather data were collected for the 29 years of trials and the six sites from MeteoSwiss (Federal Office of Meteorology and Climatology, Switzerland, consulted in 2022) using the closest weather station (Supplementary Table 1): the minimum, maximum, and average daily temperature (°C; measured 2 m above the soil surface), the daily precipitation (mm); and daily solar irradiance (W m^−2^), calculated as the average power received over 24 hours (from 0 to 24 UTC). Weather data were summed and/or averaged for each of the above-mentioned periods (*e.g.,* average of the daily average temperatures for the period from planting to harvest; sum of the daily maximum temperatures for the period from planting to harvest; etc.). This excludes the variable solar irradiance averaged over 24 hours (W m⁻²), which represents a daily mean power rather than an accumulated energy, and thus cannot be simply summed across days. We then performed two calculations: (i) it was averaged over each period to obtain the mean solar irradiance per period (W m^−2^), representing the average daily power received per square meter (averaged over 24 hours for each day then averaged over the period); and (ii) the daily values were summed for each period, then multiplied by 24 (hours) and by 0.0036 (1Wh = 0.0036 MJ) to obtain the solar radiation per period (MJ m^−2^), i.e., the cumulative energy received by the plant during that period.

The calculated climatic variables for the periods and the number of data available were recorded in the databases when available. So, we used a total of 34 climate predictors, the genotype, three measuring periods and 30 climatic variables (see list in Supplementary Table 2).

The final dataset included 29 years of variety trials conducted at six different sites and 405 varieties. The final dataset contained 10,945 records.

### 2.2) Data analysis

We used R version 4.4.1^44^ for all data preparation and statistical analyses. Packages used were the following: ggplot2^45^; tidyverse^46^; patchwork^47^; pdp^24^; vip^21^; caret^48^; party^49^; gridExtra^50^; dplyr^51^; grid^52^; doParallel^53^; sf^54^; rnaturalearth and rnaturalearthdata^55,56^; and ggrepel^57^. Detailed functions of most packages are described in sections below.

We used a two-stage analysis; first a factorial analysis quantified how genotype (G), environment (E), and their interactions (G × E) contributed to yield variability, revealing dominant factors and trends. Besides providing insights into key sources of variation, this step also supported dimensionality reduction by identifying which factors to include or replace with climatic predictors in the next step. Building on this, machine learning models (*i.e.,* decision trees and gradient boosting) were applied to explore complex, non-linear relationships between climatic variables, genotype, and grain yield.

#### 2.2.1) Factorial analysis

To assess the global effect of factors on grain yield, *i.e*., genotype (G) [N = 405]; site [N = 6], year [N = 29] and their interactions (*e.g.,* site x year = environment (E); an analysis of variance (ANOVA) was conducted separately for each decade. This approach was chosen because the full 29-year dataset was too large and complex to analyze three-way interactions in a single ANOVA. Decade-wise analysis captured temporal changes in yield variability that may diverge from national trends (Supplementary Figure 1). The ratio of sum of squares for the evaluated effect to the total sum of squares was used to obtain the percentage of the variability explained by each factor: year, site and variety, and their interactions.

#### 2.2.2) Machine learning analysis

As the effect of factors “site” and “year” on grain yield is analyzed with the factorial analysis (section 2.2.1) and these effects are global effects and can be represented through climatic variables, for the ML analysis, the factors “site” and “year” have been replaced by the 33 weather predictors described in Supplementary Table 2. In addition, a correlation matrix heatmap of numerical variables was generated in R using the cor() and corrplot functions to display Pearson correlation coefficients (Supplementary Figure 3).

In total, we had three sets of predictors: one predictor (*i.e.,* only variety as predictor), 33 predictors (only climatic variables and periods as predictors), and 34 predictors (variety and climatic variables and periods as predictors) (Supplementary Table 2), were used in two machine learning methods, a decision tree (DT) model of type “conditional inference tree”^49^ and gradient boosting (GB)^2^. Those methods were chosen because they can handle non-linear relationships and interactions while remaining compatible with post-hoc explainability tools^19,58^. DT offers clear interpretability through tree structures^21^, while GB improves prediction accuracy by aggregating multiple trees and is widely used in agronomic studies for its robustness^23^.

##### 2.2.2.1) Implementation of machine learning models (Training and Testing)

Machine learning methods were used to analyze the non-linear effects of the above-mentioned variables on grain yield. First, the dataset [N = 10945] was split randomly (80:20 split) to sort 80% of the records (*i.e.,* among all varieties, sites, year and replicates) into a training dataset [N = 8756] and 20% of the records into a test dataset used to evaluate model performance [N = 2162]. Then, the two different machine learning methods were implemented. First, for the Decision Tree we used a conditional inference tree, *i.e.,* a method that builds decision trees using statistical tests to select variables and split points, which helps avoid the variable selection bias inherent in traditional CART trees^59^ and improves interpretability^49^. We used this method because DTs are relatively simple models with high model-based interpretability^19^. Second, we implemented the GB method^23^, which combines a large number of weak learner models to achieve higher predictive accuracy. Unlike single DTs, GB models consist of hundreds to thousands of trees and require post-hoc interpretability methods to understand their behavior^19^.

The model training process included hyper-parameter tuning and a resampling using a 5-fold cross-validation to ensure robust model predictions^60^. RMSE was used to select the optimal model. To train the Decision Tree (Ctree) model, we used the train() function from the caret package in R^48^ with the method “ctree“^49^. This function fits models across a grid of tuning parameters and evaluates performance using resampling. The hyperparameter “mincriterion” was tuned over the values 0.99, 0.50, and 0.01. A total of 1,000 randomly selected samples were used with cross-validation for hyperparameter tuning, and the most consistently optimal parameter was applied to the full dataset. To train the Gradient Boosting (GBM) model, we used the same train() function with the “gbm” method. For the hyperparameter tuning the following hyperparameter grid was used: n.trees = 500, 1000, 1500, 2000, 2500; interaction.depth = 3, 6, 9, 12, 15; shrinkage = 0.1; and n.minobsinnode = 10. Final model parameters were selected based on minimum RMSE across resampling iterations. Results of the tuning process are provided in Supplementary Tables 3-4.

To test the performance of the models, predictions were calculated using the trained models with the test data set (20% of the data) using the function “predict” from the caret package in R^48^. Finally, to evaluate how well the model explains the variability of the data, the predicted values have been compared with the training values using a coefficient of determination (R-squared) based on Pearson correlation between predicted and test values. To check the effect of the factor “variety” on the model performances, the R-squared has been calculated for the three sets of predictors (see section 2.2.2) for both models with.

Finally, for both models we set seed at 123 to ensure the reproducibility of the models.

##### 2.2.2.2) Post-hoc tests to interpret ML models

IML post-hoc tests were performed to interpret machine learning analysis outcomes on both models using 33 predictors, *i.e.,* the set of data excluding the variety factor, because these tests cannot accommodate categorical variables with many levels, and our main interest was to quantify how climatic variables were treated in the models rather than varietal effects.

As described in a recent study^19^, post-hoc tests were performed to interpret the model mechanisms. This study^19^ specifically emphasizes their usefulness in ML applied to agricultural data analysis, where understanding variable interactions and the reasons behind model predictions is crucial. The tests included: 1) Permutation-based variable importance (global model-agnostic method)^20,22^, 2) pairwise interaction importance (global model-agnostic method) based on feature importance ranking measure (FIRM) approach^22^ also called “VI-based interaction statistic”, and 3) partial dependence plot (global model-agnostic method)^22,23^ (for more details see the study conducted by Ryo (2022)^19^).

The “vip” function in R^21^ was used for the “permutation-based feature importance” test to estimate the variable importance for each tested variable to predict grain yield using the R-squared as indicator. This test allows to rank the relative importance of predictors variables for prediction^19–22^.

Then, the pairwise interaction importance was tested using the function “vint()” from the “vip” package in R for both models to check key variable interactions^21,22^. This test is used to quantify the strength of two-way interaction effects affecting the model prediction^19^. Here, the variables in the top three in variable importance tests by at least one algorithm were tested. In total four variables were tested (*i.e*., “ray_p1”, “tmin_p3”, “somray_p1” and “prec_p1”) leading to six pairwise combinations.

Finally, to perform the partial dependence plot test for both models, the function “partial()” from the “pdp” package in R^24^ was used^22,23^. This test allows to visualize the modeled association between a subset of the predictors and the response taking into consideration the average effect of other predictors^19,23^.

##### 2.2.2.3) Variable importance on the most tested varieties

DT and GB models with 33 predictors (climatic variables and periods) were applied to analyze the importance of key variables. This involved comparing the top three variables identified by variable importance tests on the entire dataset to the importance of the same variables in models trained on subsets for the 15 most tested varieties, assessing their influence on popular varieties (Description of the varieties in Supplementary Table 5).

## Supporting information

Complete Supplementary Information

## Acknowledgment

We thank all colleagues and partners that have been involved in multi-environment variety testing trials over the years in different groups at Agroscope. We also thank the numerous breeders, extension services, seed producers including companies such as: “Delley Samen und Pflanzen AG”, and agricultural schools throughout Switzerland, whose collaboration and efforts in field variety trials and data collection have been essential to this research. Margot Visse-Mansiaux and Simon Treier received funding from the EU H2020 project INVITE. Josepha Schiller was funded through the Leibniz-Zentrum für Agrarlandschaftsforschung (ZALF; e. V. Müncheberg) Integrated Priority Project 2022 “CrossDiv—Co-designing smart, resilient, sustainable agricultural landscapes with cross-scale diversification”. We gratefully acknowledge the ZALF for their institutional support through scientific collaboration and for hosting Margot Visse-Mansiaux during a machine learning training.

## Author Contributions

**Margot Visse-Mansiaux:** Writing – review & editing, Writing – original draft, Visualization, Validation, Methodology, Investigation, Formal analysis, Data curation, Conceptualization;

**Masahiro Ryo:** Writing – review & editing, Visualization, Validation, Methodology, Investigation, Formal analysis;

**Amanda Burton:** Methodology, Investigation, Writing – review & editing;

**Tasin Siraj:** Methodology, Investigation, Writing – review & editing;

**Josepha Schiller:** Methodology, Investigation, Writing – review & editing.

**Simon Treier:** Data curation, Investigation, Writing – review & editing

**Didier Pellet:** Supervision, Funding acquisition, Conceptualization, Validation, Writing – review & editing

**Laura Stefan:** Writing – review & editing

**Thomas Cherico Wanger:** Supervision, Validation, Writing – review & substantial editing

**Lilia Levy Häner:** Supervision, Writing – review & editing, Validation, Data Curation, Visualization, Conceptualization.

**Juan M. Herrera:** Supervision, Funding acquisition, Writing – review & editing, Visualization, Validation, Project administration, Investigation, Data curation, Conceptualization.

