## Supplementary material for "Explainable AI shows climate impacts on wheat yields: insights from 30 years of field data": Complete Supplementary Information

**File contains:**
Supplementary Text S1.
Supplementary Figure S1 - S5
Supplementary Tables S1 - S6

### Supplementary Text

**Supplementary Text S1.** Descriptive analysis and sources of variation in grain yield across 30 years of Swiss wheat trials

#### *Descriptive analysis of grain yield variability in the dataset*

Among the 405 varieties, tested across 29 years, six sites, and three replicates, the average grain yield was 71 dt ha-1, ranging from 12.6 to 124.5 dt ha-1 [N=10945]. Grain yield followed a normal distribution (Supplementary Figure 4A). Average annual yields ranged from 53.70 dt ha-1 in 2016 [N=639] to 86.51 dt ha-1 in 2009 [N=150] (Supplementary Figure 4-D). Among the 405 varieties, average yields ranged from 43.06 dt $ha^{-1}$ to 96.29 dt $ha^{-1}$ (data not shown). Across six sites, yields averaged between 67.73 dt ha-1 at "GR" [N=636] and 74.77 dt ha-1 at "EL" [N=897] (Supplementary Figure 4-C). Supplementary Figure 4-D confirms no temporal trend in mean grain yield, ensuring data stationarity, a key assumption for ML analysis.

#### *Main factors influencing grain yield in wheat*

The ANOVA analysis showed across all decades that the most significant factor influencing grain yield was “variety,” explaining 32.47%, 34.45%, and 29.66% of yield variability for the periods 1991-2000, 2001-2010, and 2011-2020, respectively (ANOVA, $p < 0.001$; Supplementary Figure 5). While the impact of variety remained consistent over time, the importance of the “year” factor increased, surpassing the “site” factor in the second and third decades. Specifically, “site” explained 12.53% of variability from 1991-2000, while “year” accounted for 20.45% and 16.38% in the later decades. The two-way interaction “variety x site” explained 18.22%, 12.93%, and 20.33% of variability across the three decades. Similarly, the “year x site” interaction grew in importance, explaining 8.35%, 10.86%, and 15.44% over the same periods. The three-way interaction “variety x year x site” remained stable, contributing approximately 3.2%-3.8% across decades (ANOVA, $p < 0.001$; Supplementary Figure 5).

These results indicate an increasing influence of environmental factors (year x site) on grain yield over time, likely due to climate change. While global yield trends have remained steady since 1990 (Supplementary Figure 1), Swiss wheat yields have been increasingly affected by climatic variability. Our findings align with a long-term study from Kazakhstan (1972-2009), where warming and altered precipitation patterns significantly influenced grain yields in the study’s final decade[45] with climate change predicted to continue driving yield variability with some regions experiencing gains and others losses[46]. The stable three-way interaction highlights the importance of selecting varieties adapted to environmental changes and breeding in improving wheat productivity through yield parameters, disease resistance, and nutrient efficiency[47]. The observed climate differences between sites and years further justify using climatic variables to enhance yield predictions through ML analysis.

**Supplementary Figures**

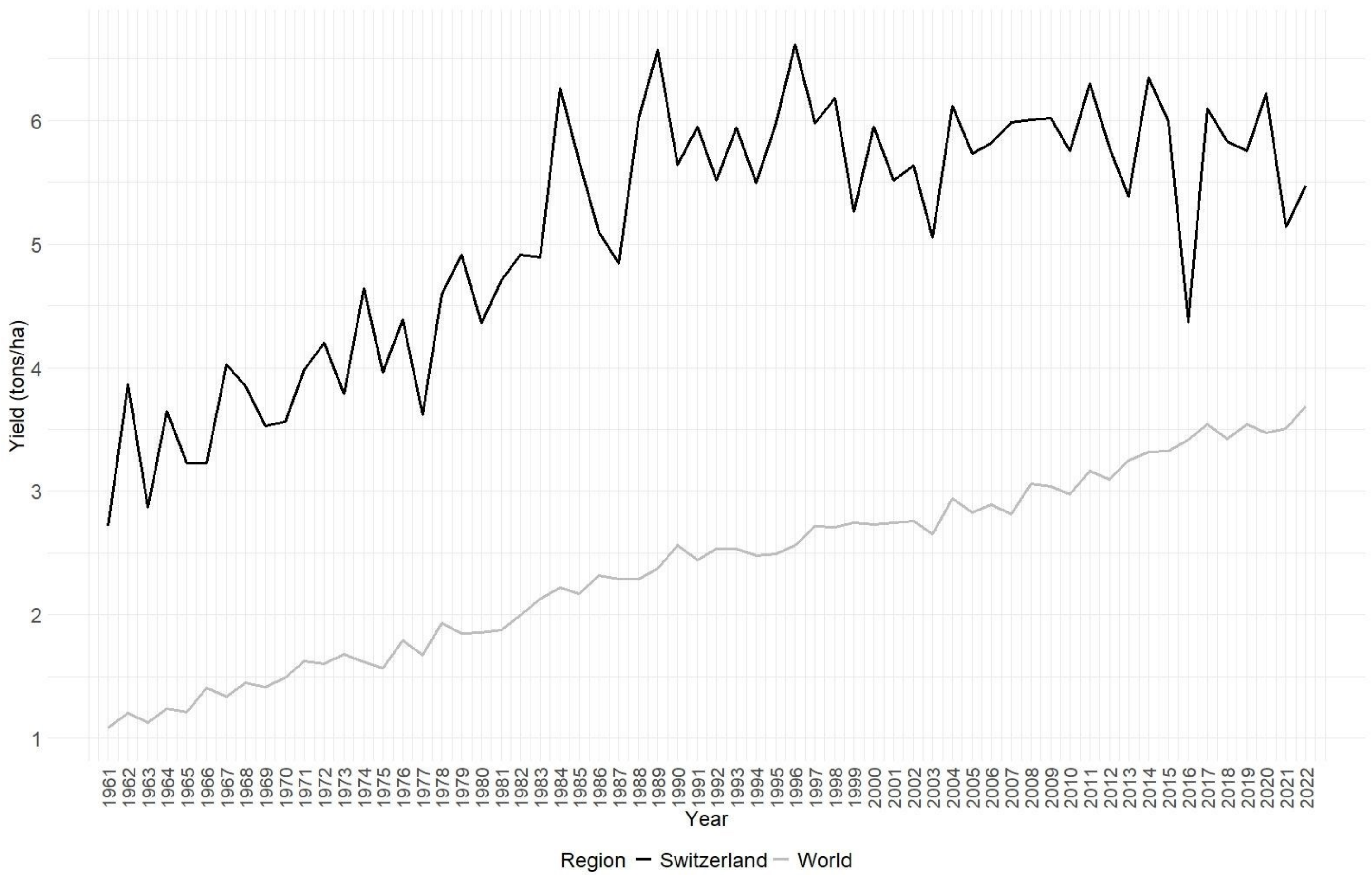


**Figure S1.** Wheat grain yield (t/ha) evolution in the world (gray) and in Switzerland (black) from 1961 to 2022 (source of data: FAOSTAT 2024).

77

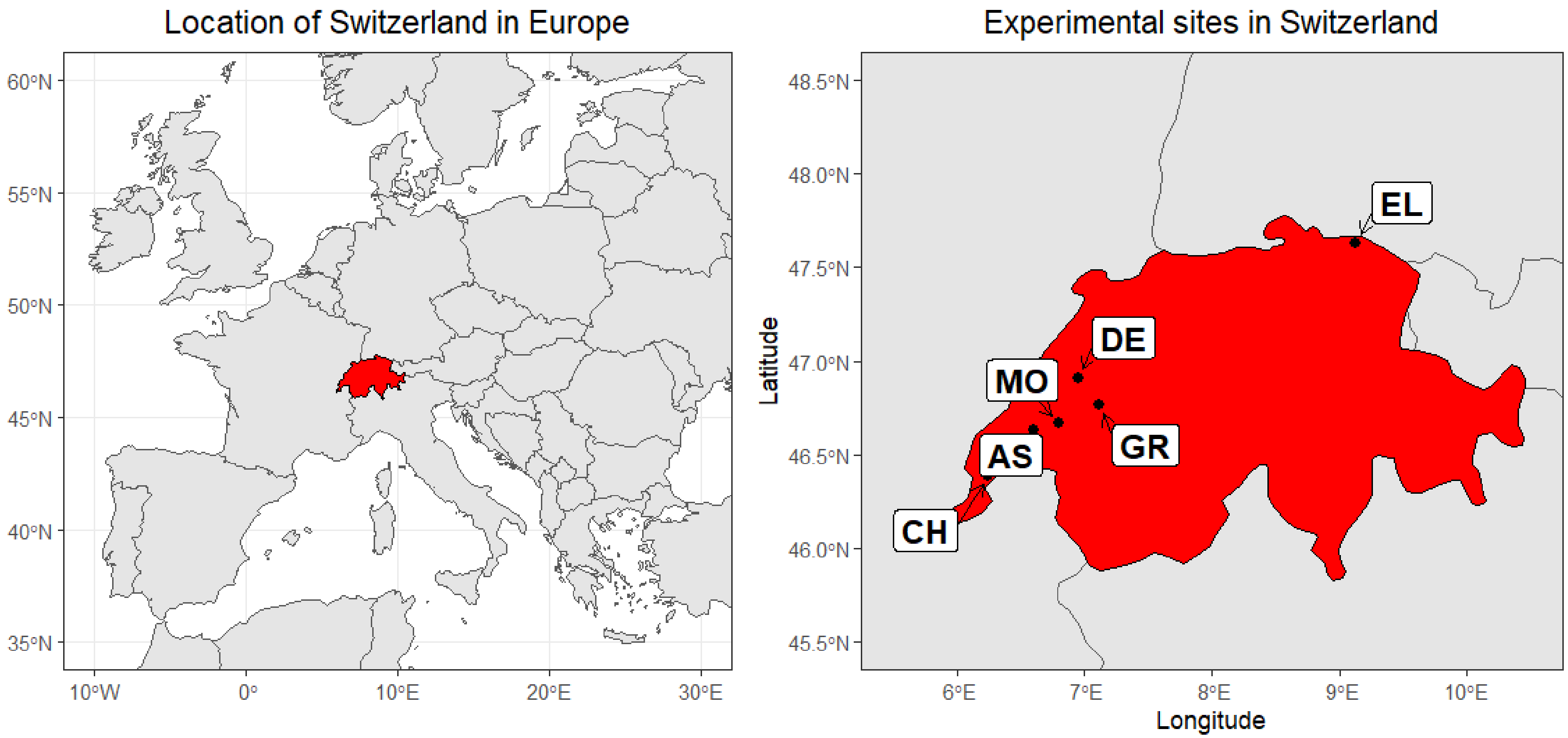


78

79 **Figure S2.** Distribution of the six sites used in the study in Switzerland.

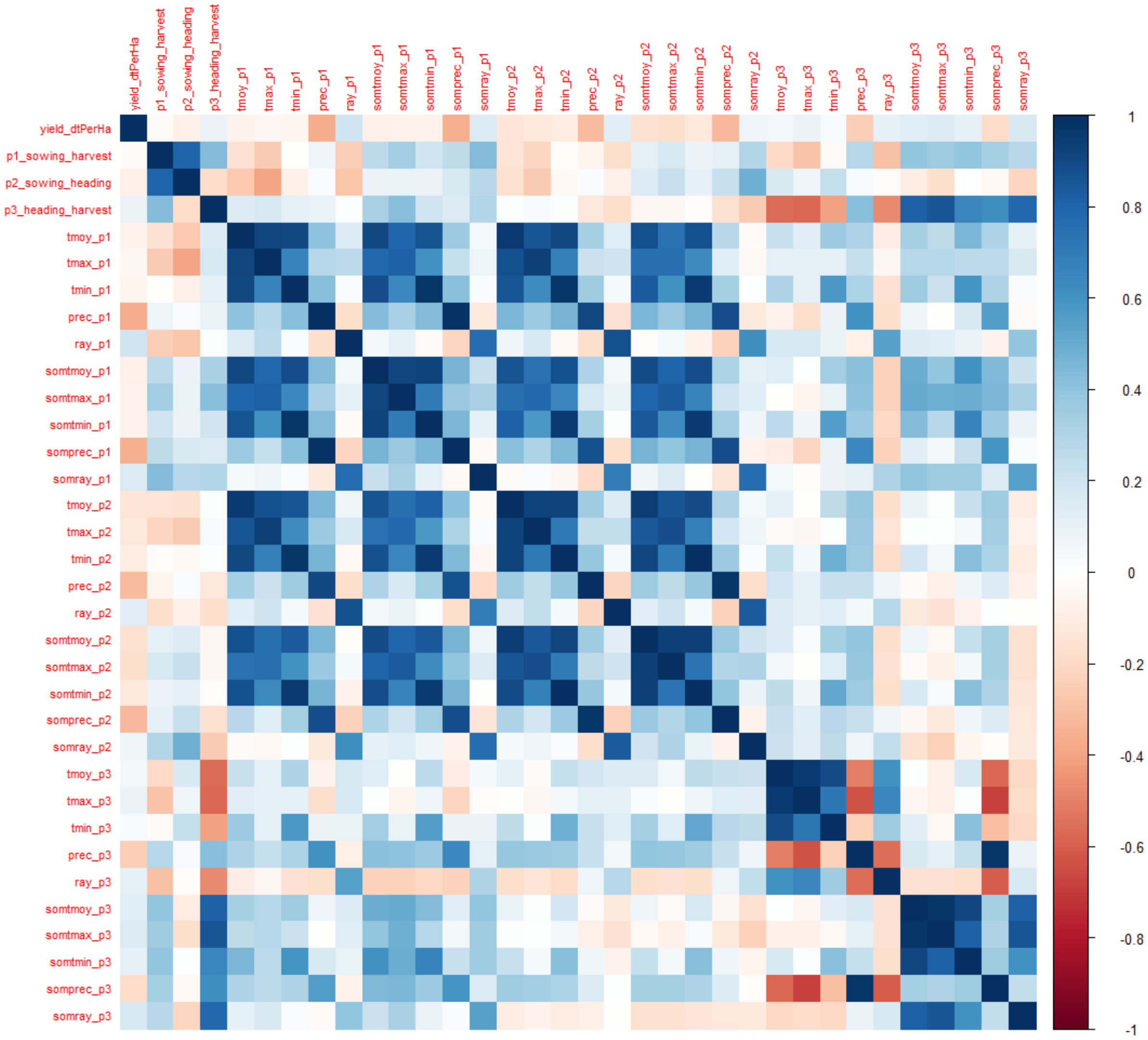


**Figure S3.** Correlation matrix heatmap created using the function “corrplot” in R applied to the correlation matrix on the numerical variables of the dataset using the “cor()” function in R showing  Pearson correlation coefficients between pairs of variables.

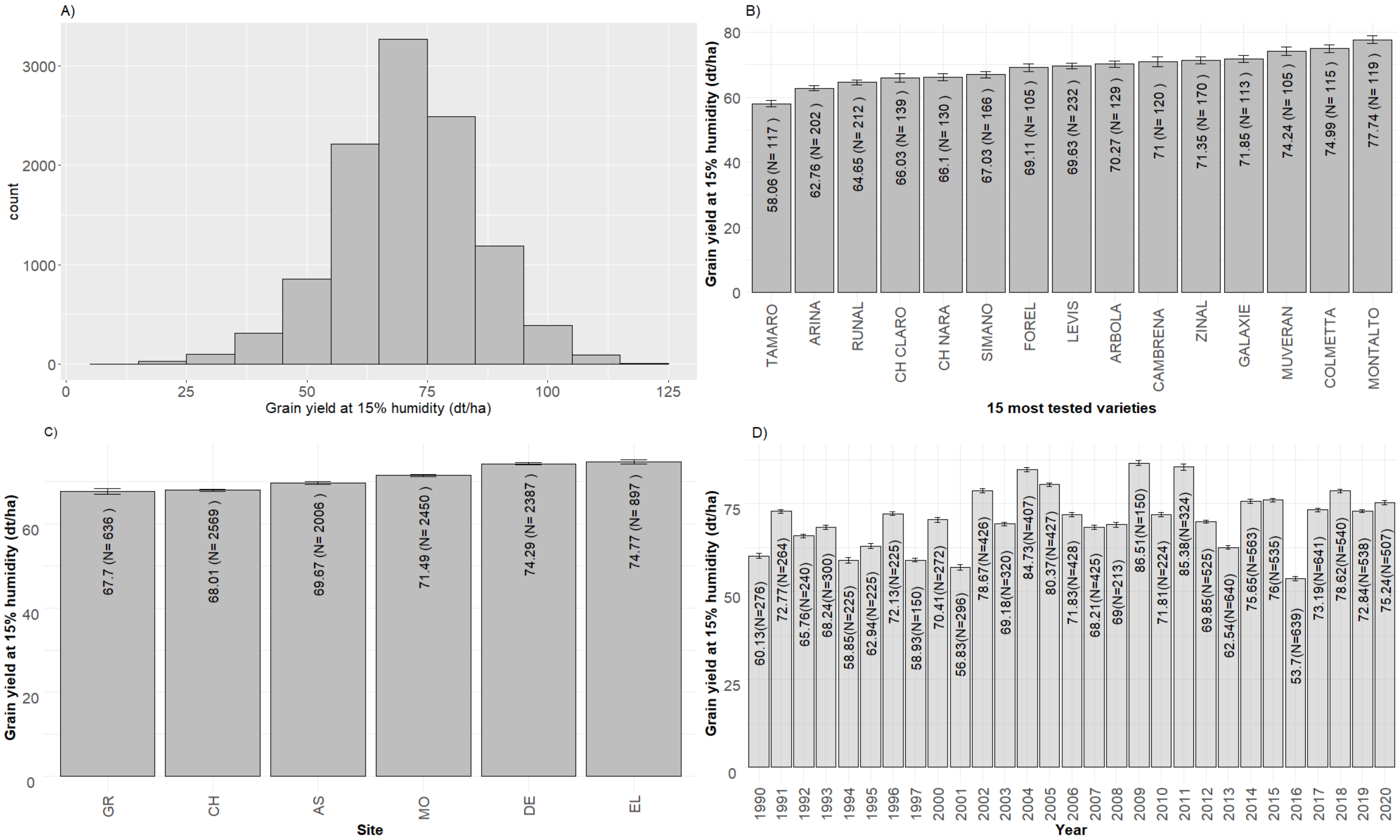


**Figure S4.** Distribution of grain yield data (A); average grain yield for the 15 most tested varieties (B); average grain yield for the six sites used in the study in Switzerland (C); average grain yield per year from 1990 to 2020 (D); average value and number of data N are indicated in the bars of the graphs (B, C, D); grain yield expressed at 15 % humidity (dt $ha^{-1}$).

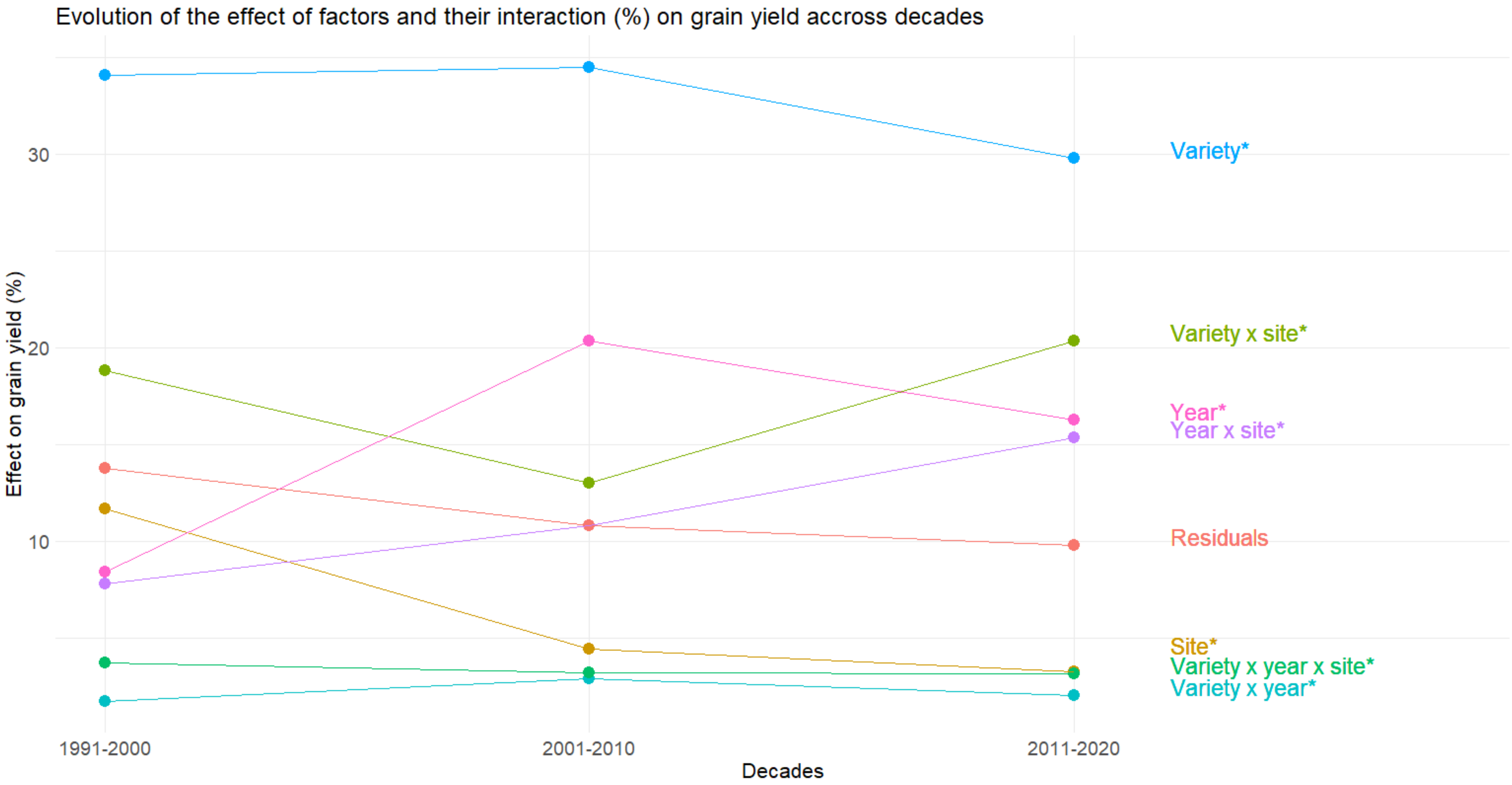


**Figure S5.** ANOVA analysis (*$p < 0.001$) to assess the effect of the main factors and their interactions on grain yield in wheat. The ratio of sum of squares for the evaluated effect to the total sum of squares was used to obtain the percentage of the variability explained by each factor.

**Supplementary Tables**

**Table S1.** Table showing information about the six sites used in the study (altitude, postal code, site names, weather station related to the site and distance from the site).

| Site | Site ID | Postal Code | Swiss canton | Weather station used for this site | Approximate distance between site and weather station [km] | Approximate elevation [m] |
|---|---|---|---|---|---|---|
| **Assens** | AS | 1042; 1049; 1040 | VD | PUY | 13 | 632 |
| **Changins** | CH | 1260 | VD | CGI | 0 to 4 | 452 |
| **Moudon** | MO | 1510 | VD | PUY | 20 | 526 |
| **Delley** | DE | 1567 | FR | NEU | 9 | 501 |
| **Grangeneuve** | GR | 1725 | FR | GRA | 0 | 641 |
| **Ellighausen** | EL | 8566 | TG | HAI | 10 | 550 |

**Table S2.** Description of the 34 predictors used for machine learning analysis (N = 10,945 per variable).

| **Predictor** | **Description of the predictor** |
|---|---|
| var | Name of the variety |
| p1_sowing_harvest | Duration (in days) from sowing to harvest (P1) |
| p2_sowing_heading | Duration (in days) from sowing to heading (P2) |
| p3_heading_harvest | Duration (in days) from heading to harvest (P3) |
| tmoy_p1 | Average of daily average temperatures (°C) during P1 |
| tmax_p1 | Average of daily maximum temperatures (°C) during P1 |
| tmin_p1 | Average of daily minimum temperatures (°C) during P1 |
| prec_p1 | Average of daily precipitation (mm) during P1 |
| ray_p1 | Average of daily solar irradiance ($W/m^2$) during P1 |
| somtmoy_p1 | Sum of daily average temperatures (°C) during P1 |
| somtmax_p1 | Sum of daily maximum temperatures (°C) during P1 |
| somtmin_p1 | Sum of daily minimum temperatures (°C) during P1 |
| somprec_p1 | Sum of daily precipitation (mm) during P1 |
| somray_p1 | Cumulative solar radiation ($MJ/m^2$) during P1 |
| tmoy_p2 | Average of daily average temperatures (°C) during P2 |
| tmax_p2 | Average of daily maximum temperatures (°C) during P2 |
| tmin_p2 | Average of daily minimum temperatures (°C) during P2 |
| prec_p2 | Average of daily precipitation (mm) during P2 |

| | |
|---|---|
| ray_p2 | Average of daily solar irradiance (W/m²) during P2 |
| somtmoy_p2 | Sum of daily average temperatures (°C) during P2 |
| somtmax_p2 | Sum of daily maximum temperatures (°C) during P2 |
| somtmin_p2 | Sum of daily minimum temperatures (°C) during P2 |
| somprec_p2 | Sum of daily precipitation (mm) during P2 |
| somray_p2 | Cumulative solar radiation (MJ/m²) during P2 |
| tmoy_p3 | Average of daily average temperatures (°C) during P3 |
| tmax_p3 | Average of daily maximum temperatures (°C) during P3 |
| tmin_p3 | Average of daily minimum temperatures (°C) during P3 |
| prec_p3 | Average of daily precipitation (mm) during P3 |
| ray_p3 | Average of daily solar irradiance (W/m²) during P3 |
| somtmoy_p3 | Sum of daily average temperatures (°C) during P3 |
| somtmax_p3 | Sum of daily maximum temperatures (°C) during P3 |
| somtmin_p3 | Sum of daily minimum temperatures (°C) during P3 |
| somprec_p3 | Sum of daily precipitation (mm) during P3 |
| somray_p3 | Cumulative solar radiation (MJ/m²) during P3 |

101

**Table S3.** Hyperparameter tuning for Ctree.

| mincriterion | RMSE | Rsquared | MAE | RMSESD | RsquaredSD | MAESD |
|---|---|---|---|---|---|---|
| 0,01 | 11,8705189 | 0,31139688 | 9,37117883 | 0,60570496 | 0,09017369 | 0,52114241 |
| 0,5 | 12,5935761 | 0,22247655 | 9,9902292 | 0,68743417 | 0,05070188 | 0,47934862 |
| 0,99 | 12,9666567 | 0,16617508 | 10,341673 | 0,85418512 | 0,06396356 | 0,81316832 |

**Table S4.** Hyper parameter tuning for GBM.

| shrinkage | interaction.depth | n.minobsinnode | n.trees | RMSE | Rsquared | MAE | RMSESD | RsquaredSD | MAESD |
|---|---|---|---|---|---|---|---|---|---|
| 0,1 | 3 | 10 | 500 | 10,1439507 | 0,49284772 | 7,82206936 | 0,51590626 | 0,04102962 | 0,51339397 |
| 0,1 | 6 | 10 | 500 | 10,5363494 | 0,46690034 | 8,0866988 | 0,45787283 | 0,03060368 | 0,48820801 |
| 0,1 | 9 | 10 | 500 | 10,8332226 | 0,44733347 | 8,37359095 | 0,56902147 | 0,04012894 | 0,61831991 |
| 0,1 | 12 | 10 | 500 | 10,7891571 | 0,45650002 | 8,28622953 | 0,50904849 | 0,04315534 | 0,56787637 |
| 0,1 | 15 | 10 | 500 | 10,9929737 | 0,43744573 | 8,43602263 | 0,50299699 | 0,0413239 | 0,61566536 |
| 0,1 | 3 | 10 | 1000 | 10,4684923 | 0,47143501 | 8,07529454 | 0,52043287 | 0,04513855 | 0,61873264 |
| 0,1 | 6 | 10 | 1000 | 10,7562261 | 0,4544779 | 8,26360407 | 0,57093773 | 0,03869311 | 0,58265052 |
| 0,1 | 9 | 10 | 1000 | 11,1191222 | 0,42978988 | 8,53548234 | 0,48922928 | 0,02959488 | 0,6074057 |
| 0,1 | 12 | 10 | 1000 | 11,1234463 | 0,43173801 | 8,54233599 | 0,52721486 | 0,04343823 | 0,58221449 |
| 0,1 | 15 | 10 | 1000 | 11,2137393 | 0,42151896 | 8,60724343 | 0,40999289 | 0,03225746 | 0,49946075 |
| 0,1 | 3 | 10 | 1500 | 10,681943 | 0,45760061 | 8,19761323 | 0,48754679 | 0,04662457 | 0,61180813 |
| 0,1 | 6 | 10 | 1500 | 10,9601299 | 0,44340779 | 8,399284 | 0,57934136 | 0,03878331 | 0,6841724 |
| 0,1 | 9 | 10 | 1500 | 11,1353382 | 0,4309042 | 8,59119029 | 0,46988895 | 0,0277168 | 0,53011785 |
| 0,1 | 12 | 10 | 1500 | 11,250696 | 0,42962945 | 8,63319434 | 0,68695685 | 0,04660027 | 0,71483775 |
| 0,1 | 15 | 10 | 1500 | 11,4040614 | 0,40757477 | 8,75478888 | 0,43173149 | 0,02441324 | 0,48829416 |
| 0,1 | 3 | 10 | 2000 | 10,8027293 | 0,45079579 | 8,33253154 | 0,54043766 | 0,0392187 | 0,6603346 |
| 0,1 | 6 | 10 | 2000 | 11,0872014 | 0,43159107 | 8,49530834 | 0,52589689 | 0,03264554 | 0,66150028 |
| 0,1 | 9 | 10 | 2000 | 11,1962368 | 0,42781313 | 8,65330808 | 0,3501086 | 0,02999523 | 0,38436311 |
| 0,1 | 12 | 10 | 2000 | 11,3587782 | 0,42342829 | 8,71389332 | 0,74924953 | 0,04835203 | 0,7140123 |
| 0,1 | 15 | 10 | 2000 | 11,3589104 | 0,41243719 | 8,72264074 | 0,38332426 | 0,03097136 | 0,47788478 |
| 0,1 | 3 | 10 | 2500 | 10,902856 | 0,44379873 | 8,41825053 | 0,4600884 | 0,0357189 | 0,66327207 |

| | | | | | | | | | |
|---|---|---|---|---|---|---|---|---|---|
| 0,1 | 6 | 10 | 2500 | 11,15 39577 | 0,429 73986 | 8,526 81604 | 0,477 70088 | 0,028 39075 | 0,603 14818 |
| 0,1 | 9 | 10 | 2500 | 11,27 19195 | 0,424 42315 | 8,677 76725 | 0,433 16427 | 0,027 14134 | 0,407 43039 |
| 0,1 | 12 | 10 | 2500 | 11,43 95054 | 0,420 71323 | 8,767 11106 | 0,734 70724 | 0,050 23686 | 0,640 07195 |
| 0,1 | 15 | 10 | 2500 | 11,50 83455 | 0,402 0799 | 8,844 31145 | 0,380 97743 | 0,040 95264 | 0,440 30087 |

**Table S5.** Description of the 15 most tested varieties (sources: *EUPVP – COMMON CATALOGUE, https://ec.europa.eu/food/plant-variety-portal/ (consulted on October 2024); **Swiss variety list, the most recent list containing the variety was selected, lists obtained from the website of Swissgranum https://www.swissgranum.ch/fr/directives/varietes; or the Agroscope repository https://ira.agroscope.ch/ (Consulted on October 2024) or internal files for old lists).

| Variety | Country* | Maintainer (national ID)* | Breeder's Ref* | Status* | Quality class in Switzerland**[1] | Year of registration in the Swiss catalogue** | Yield potential (Extenso production)**[2] | Yield potential (PER production)**[2] | Last recommended list with the variety for the year's harvest**: |
|---|---|---|---|---|---|---|---|---|---|
| Tamaro | CH / FR | DSP (41) for CH / DELLEY SEMENCES ET PLANTES SA (11444) for FR | none | Surrendered | TOP | 1992 | Ø | ? | 2004 |
| Arina | CH | DSP (41) | CH 72077 | registered | I | 1981 | - | - | 2024 |
| Runal | CH / FR | DSP (41) for CH / DELLEY SEMENCES ET PLANTES SA (11444) for FR | CH 75269 (CH) / none (FR) | Registered (CH) / surrendered (FR) | TOP | 1995 | - | - - | 2024 |
| CH Claro | CH | DSP (41) | CH 111.12754 | registered | TOP | 2009 | - | + | 2023 |
| CH Nara | CH | DSP (41) | CH 111.13197 | registered | TOP | 2010 | + | - | 2024 |
| Simano | CH | DSP (41) | CH 111.13726 | registered | I | 2012 | + | Ø | 2023 |
| Forel | CH | DSP (41) | CH 111.12943 | registered | I | 2008 | + | + | 2024 |
| Levis | CH / FR / IT | DSP (41) for CH / DELLEY SEMENCES ET PLANTES SA (11444) for FR / DELLEY SEMENCES ET PLANTES SA (2004) for IT | CH 75802 (CH) / none (FR) / none (IT) | Registered (CH) / surrendered (FR) / surrendered (IT) | II | 1997 | ++ | + | 2024 |
| Arbola | CH | DSP (41) | none | Surrendered | Biscuit | 1994 | + | ++ | 2005 |
| Cambrena | CH | DSP (41) | CH 194.10119 | registered | Biscuit | 2011 | +(+) | ++ | 2022 |
| Zinal | CH | none | none | Surrendered | I | 2003 | + | Ø | 2018 |
| Galaxie | CH / FR | R2N (116) for CH / R 2N (S12452) for FR | CH 60975 (CH) / none (FR) | Registered (CH) / surrendered (FR) | II | 1991 | + | ++ | 2013 |
| Muveran | CH | none | none | Surrendered | Biscuit | 2004 | + | + | 2011 |
| Colmetta | CH | none | none | Surrendered | NA | NA | NA | NA | NA |
| Montalto | CH | DSP (41) | CH 111.14316 | registered | II | 2016 | ++++ | ++(+) | 2022 |

Legend: [1]The quality class of winter wheat is defined using an overall quality index and thresholds for wet gluten content, with requirements as follows: TOP >130 points, ≥31% gluten, agronomic index >95; Class I >110–130, ≥29%, >103; Class II >95–110, ≥27%, >110; Class III >80–95, no %, >115; Feed ≤80, no %, >120; Biscuit class has specific criteria, with agronomic index >110; values measured in PER trials, thresholds may vary depending on the year's conditions. [2]: ++++ = excellent; +++ = very good; ++ = good; + = fair to good; Ø = average; - = moderate to low; -- = low; --- = very low; ? = no information. NA= no information because the variety was not in the variety lists.

**Table S6.** Relevant values from the decision tree and gradient boosting best models (*i.e.*, with the smallest RMSE) for each dataset (*i.e.*, 1, 33, or 34 predictors). The DT and GB models were performed on the train dataset [N = 8756] using 5-cross resampling with different tested sample sizes (*i.e.*, 7005, 7005, 7005, 7005 and 7004).

| Model | Number of predictors | Description of predictors | RMSE | R2 | MAE | Mincriterion | n.trees | shrinkage | n.minobsinnode | interaction.depth |
|---|---|---|---|---|---|---|---|---|---|---|
| **Decision tree** | 1 | Variety | 14.19 | 0.01 | 11.06 | 0.99 | None | None | None | None |
| | 33 | Climatic variables and periods | 9.29 | 0.57 | 7.25 | 0.01 | None | None | None | None |
| | 34 | Variety, climatic variables and periods | 10.3 | 0.48 | 8.04 | 0.01 | None | None | None | None |
| **Gradient boosting** | 1 | Variety | 12.81 | 0.19 | 9.99 | None | 2500 | 0.1 | 10 | 12 |
| | 33 | Climatic variables and periods | 8.79 | 0.62 | 6.87 | None | 500 | 0.1 | 10 | 6 |
| | 34 | Variety, climatic variables and periods | 7.4 | 0.73 | 5.7 | None | 2500 | 0.1 | 10 | 3 |
